# Orthogonality of shell proteins across BMC subclasses in cyanobacteria

**DOI:** 10.1101/2024.03.19.585794

**Authors:** Joshua S. MacCready, Matthew E. Dwyer, Cheryl A. Kerfeld, Daniel C. Ducat

## Abstract

Bacterial microcompartments (BMC) are protein-based organelles broadly distributed across all bacterial phyla and subclassified into ≥60 functional variants. Despite their evolutionary and metabolic diversity, shell proteins that structurally compose the BMC surface are closely related across BMC classes. Herein, we sought to identify molecular and physiological features that could promote independent operation of more than one BMC type within the same cell by reducing inter-organelle cross-talk of shell proteins. We heterologously expressed shell proteins from the structurally well-defined BMC of *Haliangium ochraceum* (HO) within *Synechococcus elongatus* PCC 7942, a model cyanobacterium containing the β-carboxysome. We find considerable cross-reactivity of the HO hexameric shell protein (HO BMC-H) with components of the β-carboxysome; HO BMC-H can integrate into carboxysomes, disrupt its ultrastructural organization, and impair its associated CO_2_ fixation reactions. *S. elongatus* is unable to maintain the integrity of the β-carboxysome over time when HO BMC-H is expressed in the absence of one or more of three broad strategies that act to increase the orthogonality between HO and carboxysome BMC shell proteins: i) reduced expression of promiscuous shell proteins; ii) sequestration of free HO BMC-H proteins via co-expression of other members of the same HO shell protein class, or; iii) heterologous expression of BMC positional system proteins McdAB (<u>M</u>aintenance of <u>c</u>arboxysome <u>d</u>istribution AB), revealing a putative moonlighting function of the McdAB protein family. Our results have implications for bacteria that encode more than one BMC within their genome and may have translational implications for the use of engineered BMCs for biotechnological applications.

## Introduction

Bacterial microcompartments (BMCs) are protein-based organelles that are increasingly recognized to increase the efficacy of sensitive enzymatic reactions by creating specialized subcellular environments favorable to a given metabolic pathway **(Reviewed in: Kerfeld et al., 2018; Kirst and Kerfeld, 2019)**. Recent bioinformatic surveys have highlighted the ubiquity of this bacterial compartmentalization strategy, identifying at least 60 different classes of BMC’s that encapsulate distinct metabolic pathways encoded across at least 45 bacterial phyla **(Axen et al., 2014; Sutter et al., 2021)**. While the phylogenetic distribution of BMCs is wide and the metabolism contained within is thought to be very diverse, BMCs share several common features. For example, BMCs appear to improve flux by concentrating key intermediates that are otherwise reactive or prone to toxic/wasteful side reactions that can reduce metabolic efficiency. The model anabolic BMC is the carboxysome, encoded in all cyanobacteria and many chemoautotrophic proteobacteria, which facilitates carbon fixation by concentrating CO_2_ near enzymatic active sites of Ribulose-1,5-bisphosphate carboxylase/oxygenase (Rubisco) in the lumen **(Rae et al., 2013)**. Rubisco has dual enzymatic activities of fixing either CO_2_ (carboxylation) or O_2_ (oxygenation), the latter of which can be an energetically expensive side reaction requiring photorespiratory bypasses. Due to the selective permeability of the protein shell that defines the carboxysome, bicarbonate can traverse the BMC shell and be converted by carbonic anhydrase to generate a high internal CO_2_ environment favoring Rubisco reactions that drives flux towards the Calvin–Benson–Bassham cycle and away from the wasteful process of photorespiration **(Reviewed in: Rae et al., 2013; Turmo et al., 2017)**. Analogously, the best-studied catabolic BMCs act to concentrate reactive pathway intermediates (*e.g*., aldehydes) near the enzymes involved in processing them, thereby preventing their entrance into the cytosol where many toxic side-reactions can result. The significant metabolic enhancements BMCs enable confer measurable growth advantage to the organisms that encode them under physiologically-relevant environments **(Prentice, 2021; Stewart et al., 2021)**.

Another similarity that transcends different classes of BMCs are the shell proteins that assemble into the selectively-permeable outer barrier of the compartment. Shell proteins are structurally and evolutionarily related, containing 3 broad classes: hexamers (BMC-H; Pfam00936), pentamers (BMC-P; Pfam03319), and pseudohexameric trimers (BMC-T; tandem repeats of the Pfam00936 domain) **(Kerfeld et al., 2005; Tanaka et al., 2008; Klein et al., 2009)**. These proteins form the facets (BMC-H and BMC-T) or vertices (BMC-P) of the polyhedral ∼40-200 nm shape adopted by all BMC’s **(Figure 1A)**. Shell proteins share homology, and while surface electrostatics and features of the pores that mediate small substrate transport across the shell can vary, the overall protein fold is highly conserved across evolutionarily distant BMC classes **(Chowdhury et al., 2015; Sommer et al., 2017; Melnicki et al., 2021)**. While the genomes of existing laboratory models contain only one class of BMC, recent metagenomic analyses have revealed that many bacterial species encode 2 or more BMC classes (>20 % of BMC-containing bacteria) **(Sutter et al., 2021)**. This is illustrated by *Maledivibacter halophilus*, an anaerobic chemoorganotroph, which encodes 6 different BMC classes in its genome **(Sutter et al., 2021)**. Given that BMCs are horizontally transferred among microbes, BMC incompatibility could exist, whereby organisms are selected for or against depending on whether the BMC operon that has been transferred interferes with assembly or functionality of any prior BMC types.

**Figure 1:**
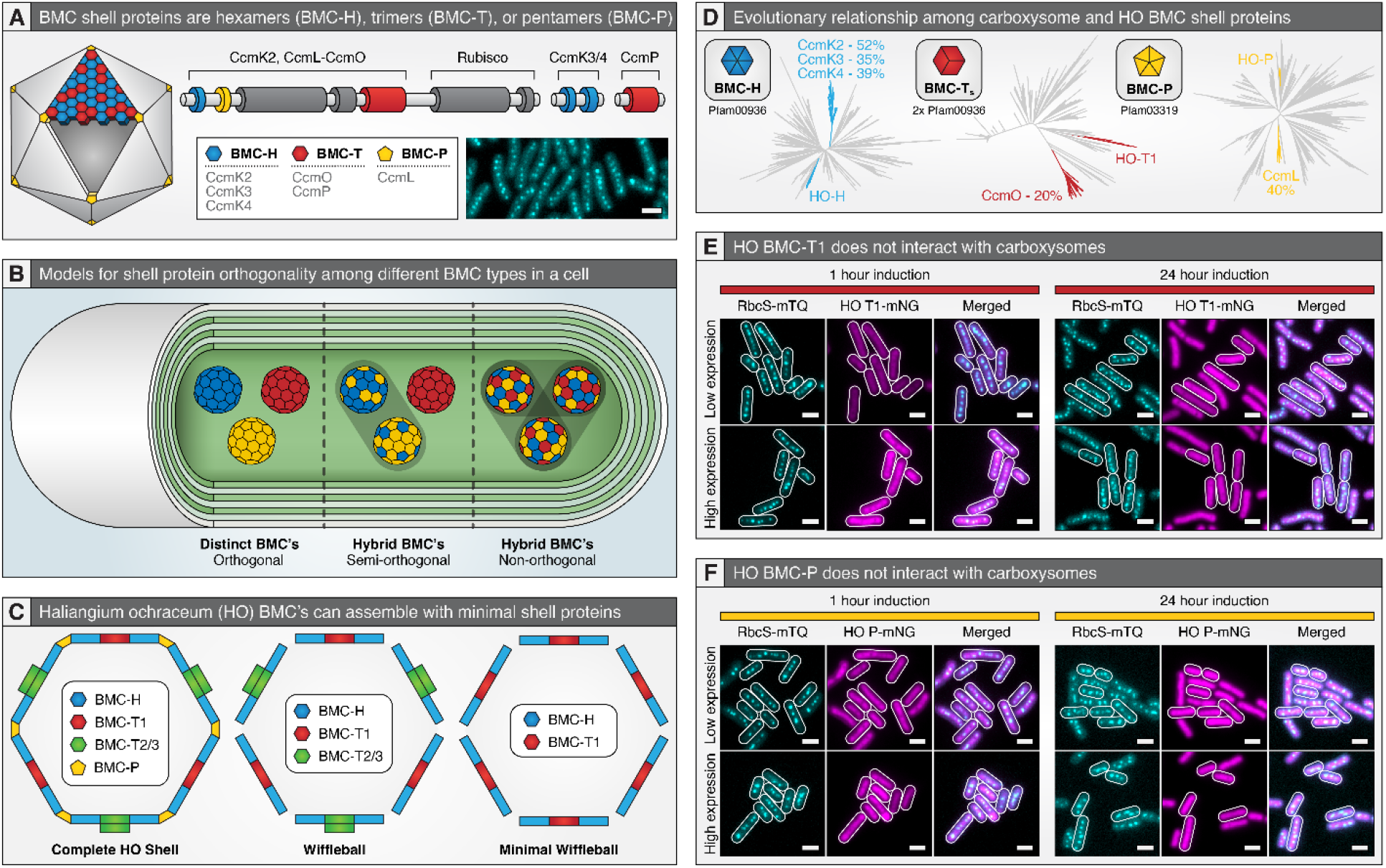
BMC shells are composed from three protein families. (A) The main carboxysome operon of S. elongatus 7942 encodes for BMC-H (blue), BMC-T (red), and BMC-P (yellow) shell proteins, with additional BMC-H and BMC-T proteins found at additional loci. Inset: Carboxysomes are visualized by expressing an additional copy of RbcS-mTQ. (B) Illustration demonstrating possible shell orthogonality outcomes in cells co-expressing more than one BMC type. (C) Cartoon schematic of minimal shell protein assemblies previously reported by expression of 2 to 5 HO BMC shell subunits. (D) Phylogenetic inference of selected BMC-H, BMC-T, and BMC-P shell proteins highlighting relatedness of specific shell proteins utilized in this study. For a more comprehensive phylogenetic analysis, see ([Melnicki et al., 2021; Sutter et al., 2021)]. (D) Phylogenetic inference of BMC-H, BMC-T, and BMC-P proteins to demonstrate relatedness between carboxysome and HO BMC shell proteins. (E) Signal from the expression of HO BMC-T1-mNG and (F) HO BMC-P-mNG does not colocalize with the carboxysome reporter Rbcs-mTQ. Scale bars = 2 µm.

The high conservation of key structural components of BMCs alongside the prevalence of organisms with multiple BMC classes raises many fundamental questions regarding BMC evolution and orthogonality, including: (i) Are shell proteins interchangeable? Or do shell proteins exhibit orthogonality, meaning they can only interact with other shell proteins within their specific BMC subtype?; (ii) If shell proteins do interact across BMC classes, does cross-class integration disrupt important features of BMC function (*e.g*., selective permeability)?; and iii) Do shell proteins have inherent features that restrict their capacity to interact with distantly-related shells of a different BMC, or are they granted specificity by interaction with other components of the BMC (*i.e*., interactions with “core” components) **(Figure 1B)**? Because bacteria that naturally encode more than one BMC class have not been well-studied, it is an open question if more than one BMC type can be functionally maintained at the same time within a cell.

Minimal BMC systems have been identified that are relatively simple in comparison to the complexity of the best-studied metabolosome systems. One well-characterized example of a BMC that can be assembled from a limited set of shell proteins is derived from the myxobacterium *Haliangium ochraceum* (HO BMC), composed of just a single BMC-H, 3 BMC-T, and 1 BMC-P that can self-assemble *in vitro* as an empty shell with a solved structure **(Lassila et al., 2014; Sutter et al., 2017) (Figure 1C)**. This stands in contrast to the multi-gene BMC arrays of the 1,2-propanediol utilization (PDU), ethanolamine utilization (EUT), and glycyl radical enzyme-associated (GRM) BMCs, which possess anywhere from 6 to 8 shell proteins in their respective model species **(Sutter et al., 2021)**. Furthermore, the HO BMC shell proteins can self-assemble with an even more streamlined set of shell proteins *in vivo* and *in vitro*, such as the “wiffle ball” (**Kirst et al., 2022)** composed of a single BMC-H and BMC-T1 **(Figure 1C) (Hagen et al., 2018ab; Ferlez et al., 2019; Sutter et al., 2019)**. By virtue of its simplicity, the minimal HO BMC system represents a good candidate to explore questions related to orthogonality posed above.

Here, we express shell proteins of the HO minimal BMC in the model rod-shaped cyanobacterium *Synechococcus elongatus* PCC 7942 (*S. elongatus*). We found that while HO BMC-T1 and HO BMC-P exhibit no evidence of co-localization or interference with carboxysomes, HO BMC-H displayed considerable interchangeability between different classes of shell protein assemblies. While HO BMC-H initially co-localized with other HO shell proteins and appears to form spherical structures by transmission electron microscopy (TEM), when expressed under higher induction or over a longer duration, HO BMC-H colocalized with carboxysomes and severely disrupted native carboxysome morphology and function. We explore 3 different strategies that each can partially act to insulate the carboxysome and HO shell assemblies from one another – results that have implications for how bacteria may maintain more than one BMC class that operate simultaneously, yet independently.

## Results and Discussion

### HO BMC-T1 and BMC-P proteins are orthogonal to the cyanobacterial carboxysome

We sought to introduce a heterologous minimal shell BMC system from the myxobacterium *Haliangium ochraceum* (HO BMC) into the model rod-shaped cyanobacterium *S. elongatus*. Cyanobacteria were selected not only because they possess the model carbon fixing BMC, the β-carboxysome, but also because carboxysomes are essential to cyanobacterial carbon fixation under atmospheric concentrations of CO_2_, thus disruptions to carboxysome function result in a readily-identified growth phenotype (high CO_2_ requirement; **Cameron et al., 2013; Rae et al., 2013)**. Carboxysome components in *S. elongatus* are largely encoded within a single operon, *ccmK2LMNO*, with additional accessory shell proteins CcmK3/4 and CcmP encoded in satellite loci **(Figure 1A)**. HO BMCs have homologous shell proteins to the carboxysome of *S. elongatus*, including: (i) HO BMC-H, which shares ∼ 52% pairwise identity to CcmK2, 35% pairwise identity to CcmK3, and 39% pairwise identity to CcmK4, (ii) HO BMC-T1, which share ∼ 20% pairwise identity to CcmO, and (iii) HO BMC-P, which shares ∼ 40% pairwise identity to CcmL **(Figure 1D)**. Therefore, the relative evolutionary distance of the HO BMC shell proteins - both by sequence identity and a more comprehensive recent phylogenetic analysis **(Melnicki et al., 2021)** - suggest that they might be good candidates to act orthogonally to the carboxysome.

We began by expressing individual HO shell proteins (HO BMC-H, HO BMC-T1, and HO BMC-P) and assaying whether they localized to carboxysomes and/or perturbed carboxysome morphology, function, or positioning. To monitor the localization and morphology of carboxysomes, we created a carboxysome reporter background by genomically-integrating an additional copy of the small subunit of Rubisco (RbcS) fused at the c-terminus to mTurquoise2 (mTQ) under a second copy of the native *rbcS* promoter, as has been characterized previously **(MacCready et al., 2018; Hakim et al., 2021; Rillema et al., 2021)**. Simultaneously, HO BMC-H, HO BMC-T1, or HO BMC-P fluorescently fused at the c-terminus to mNeonGreen (mNG) and expressed using a synthetic riboswitch was inserted upstream of the carboxysome reporter.

We found that heterologous HO BMC-T1-mNG and HO BMC-P-mNG exhibited orthogonality to carboxysomes regardless of the degree or duration of protein expression. HO BMC-T1-mNG and HO BMC-P-mNG shell proteins did not co-localize with carboxysomes at any time following induction, retaining a diffuse and cytosolic localization pattern **(Figure 1EF)**. Moreover, consistent with previous *in vitro* characterizations **(Lassila et al., 2014)**, expression of HO BMC-T1 or HO BMC-P did not show evidence of self-assembly into higher-order structures discernible by light microscopy, such as sheets or filaments. This diffuse pattern of HO BMC-T1 and HO BMC-P was consistent even at the highest expression levels allowed under our inducible system.

### HO BMC-H proteins can localize to the cyanobacterial carboxysome

In contrast to other HO shell proteins, we found that HO BMC-H-mNG exhibited the capacity to form foci-like puncta in the cytosol within the first hour of expression, regardless of expression intensity **(Supplementary Figure 1A)**, a propensity that increased when visualized after 4 hours of expression **(Figure 2A – left panel)**. This observation could be suggestive that HO BMC-H is oligomerizing to form sheets or other small structures, as previously reported for this shell protein *in vivo* or *in vitro* **(Lassila et al., 2014; Sutter et al., 2016)**. In addition to this separate cytosolic pool of HO BMC-H-mNG foci, a subfraction of the signal was strongly concentrated near 1 or more carboxysomes, as visualized by the RbcS-mTQ reporter **(Figure 2A)**. At 1-hour post-induction, limited colocalization was observed **(Supplementary Figure 1A)**, we observed that HO BMC-H-mNG signal was concentrated near ∼ 28% of carboxysomes at 4 hours of expression **(Figure 2A – left panel)** and ∼ 85% of carboxysomes after 24 hours of expression **(Figure 2A – right panel)**. Given the doubling-time of ∼8 hours for *S. elongatus* under our experimental conditions, a possible interpretation is that HO BMC-H may only be capable of association with carboxysomes while they are newly forming, but that excess shell protein does not efficiently interact with previously formed carboxysomes. An alternative possibility is that higher-order HO BMC-H assemblies may stochastically nucleate near or on the surface of carboxysomes that cannot be separately resolved at the limits of light microscopy.

**Figure 2:**
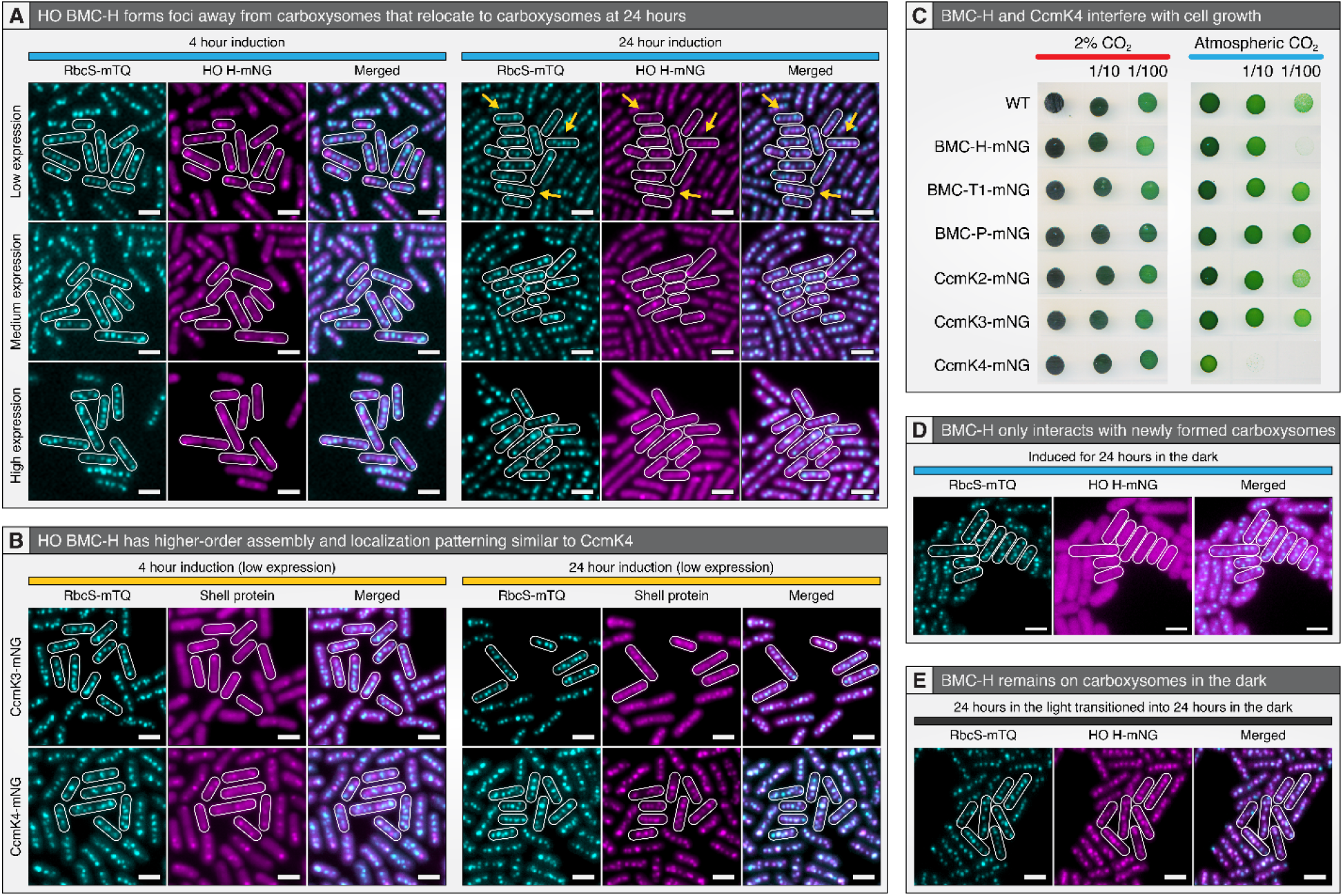
HO BMC-H co-localizes with carboxysomes and its expression perturbs cellular growth on atmospheric CO_2_. (A) HO BMC-H-mNG forms puncta at early timepoints that are not always co-localized with the carboxysome reporter yet relocates to carboxysomes at 24 hours (yellow arrows). (B) Additional fluorescent reporter copies of the endogenous carboxysome shell proteins CcmK3-mNG and CcmK4-mNG demonstrate similar localization patterns to HO BMC-H-mNG. (C) Serial diluted spot plate assay with HO and carboxysome shell proteins at 2% and atmospheric CO_2_ levels. (D) HO BMC-H does not form punctate structures or interact with carboxysomes when expressed for up to 24 hours in cells incubated in the dark. (E) HO BMC-H that colocalized to carboxysomes after 24 hours induction in the light does not relocate when cells are transitioned to the dark for 24 hours. Scale bars = 2 µm.

In an attempt to disambiguate whether HO BMC-H-mNG was interacting with the carboxysome surface or being integrated with native shell proteins during carboxysome facet formation, we examined the kinetics of native carboxysome shell proteins localization using the same inducible reporter approach. The major carboxysome shell protein CcmK2, and minor paralogs CcmK3 and CcmK4, were individually fluorescently tagged at their C-terminus and expressed from the same inducible promoter as HO BMC-H above (*i.e*., CcmK2-mNG, CcmK3-mNG, or CcmK4-mNG). We observed that the minor shell proteins CcmK3 and CcmK4 exhibited localization kinetics similar to that of HO BMC-H. CcmK4-mNG behaved the most similarly to HO BMC-H, with higher-order assembly into puncta at early timepoints that colocalized to a subset of carboxysomes at 4 hours and nearly all carboxysomes at 24 hours **(Figure 2B)**. CcmK3-mNG remained diffuse at 1 hour of expression **(Supplemental figure S2A)**. The results show that HO BMC-H has a similar pattern of localization over time with the native minor shell proteins CcmK3 and CcmK4, providing supporting evidence that HO BMC-H might be directly integrating into the shell of carboxysomes. Although CcmK3 associates with carboxysomes with slower kinetics (4 hours vs. 1 hour), the recent determination that CcmK3 cannot form homohexamers and instead can only form heterohexamers with CcmK4 **(Sommer et al., 2019)** may explain this slower association. Alternatively, we observed that CcmK2-mNG was largely diffuse throughout most time points, but eventually colocalized with carboxysomes after 24 hours. In many instances, it appeared that a high background signal of unbound CcmK2-mNG partially masked the colocalization signal, even at low expression **(Supplemental figure S2A)**. This larger background signal may be the result of a number of possible effects, including: (i) less efficient integration of tagged CcmK2-mNG than endogenous CcmK2 **(Sun et al., 2019)**; (ii) saturation of the possible CcmK2 binding sites in carboxysomes, and/or; (iii) a higher basal level of CcmK2 protein levels in competition to bind to carboxysomes **(Cameron et al., 2013)**.

Expression of different shell protein reporters had a varied impact on cell growth at ambient CO_2_ levels. We observed a slow growth phenotype at atmospheric CO_2_ for HO BMC-H-mNG and CcmK4-mNG in comparison to WT and other reporters; the slow growth phenotype of CcmK4-mNG was most severe **(Figure 2C)**. This decrease in growth was alleviated when the same strains were grown under an enriched CO_2_ headspace (2%). This suggests over-expression of both HO BMC-H and CcmK4 could partially compromise the functionality of the carboxysome, without fully disrupting the cyanobacterial carbon concentrating mechanism.

To further evaluate the hypothesis that HO BMC-H-mNG might be integrating directly into the native carboxysome shell, rather than aggregating near the compartment, we manipulated illumination to modulate the rate of carboxysome formation. Since *S. elongatus* is an obligate photoautotroph, cells do not divide in the absence of light and carboxysome biogenesis is halted. Interestingly, we neither observed the formation of distinct cytosolic foci or colocalization of HO BMC-H-mNG with carboxysomes when it was expressed while cells were cultivated in the dark for 24 hours **(Figure 2D)**. Finally, we evaluated the possibility that BMC-H-mNG can only interact at the surface of carboxysomes, but that this association is pH-dependent. Carboxysomes are hypothesized to maintain a pH gradient driven by internal metabolism during the day that acidifies the cytosol in their vicinity **(Menon et al., 2010; Whitehead et al., 2014; Mangan et al., 2016; Long et al., 2021; MacCready and Vecchiarelli, 2021)**. Since this pH gradient collapses in the absence of light **(Mangan et al., 2016)**, we tested whether BMC-H-mNG already colocalized to carboxysomes relocated to the cytosol after 4 or 24 hours of dark exposure. However, HO BMC-H-mNG did not dissociate from carboxysomes after exposure to 24 hours of darkness **(Figure 2E and Supplemental figure S2B)**.

Taken together, our data suggests that HO BMC-H is likely integrating into carboxysome shells given the similarities between HO BMC-H-mNG localization patterns with that of the native minor shell proteins (CcmK3 and CcmK4) and growth phenotypes at atmospheric CO_2_. The interaction of HO BMC-H with carboxysomes is similar to results found for other BMC-H family proteins, such as those from CsoS1A from α-carboxysomes and PduA from the propanediol utilization BMC, which have also been suggested to colocalize with *S. elongatus* carboxysomes **(Cai et al., 2015; Fang et al., 2018; Zedler et al., 2023)**. Moreover, given that HO BMC-H-mNG co-localized to ∼28% of carboxysomes at 4 hours of expression and ∼85% of carboxysomes at 24 hours of expression, this further suggests that HO BMC-H is integrating preferentially into shells of newly formed carboxysomes since these numbers correlate closely to the theoretical quantity of newly synthesized carboxysomes from the time of induction (25% at 4 hours and 87.5% at 24 hours, respectively).

### Co-expression of HO BMC-T1 and HO BMC-H forms transiently-separate foci that don’t maintain orthogonality

We next asked whether HO wiffle balls could form within living *S. elongatus*, or if the apparent cross-reactivity of some HO shell proteins would preclude this possibility. The most basic structure of the HO BMC shell is the minimal wiffle ball, which robustly assembles *in vitro* and is composed of just HO BMC-H and HO BMC-T1, lacking capping of the vertices by HO BMC-P **(Figure 1C) (Lassila et al., 2014)**. We therefore first inserted untagged HO BMC-H into our HO BMC-T1-mNG strain containing the carboxysome reporter.

The localization pattern of HO BMC-T1 was significantly altered when co-expressed with HO BMC-H **(Figure 3A)**, relative to when it was expressed in isolation **(Figure 1E)**. HO BMC-T1-mNG signal relocalized from a diffuse pattern **(Figure 1E)** into small puncta that did not colocalize with carboxysomes when co-expressed with HO BMC-H **(Figure 3A)**. The kinetics of formation for these separate puncta could be tuned as a function of the expression level: at high expression, they formed at 1 hour, at medium expression, by 4 hours, and only by 24 hours at low expression. These puncta exhibited a rapid rate of diffusion within the bacterial cytosol that was readily distinct from the diffusion of carboxysome puncta, which are tethered to the bacterial nucleoid via the maintenance of carboxysome distribution A and B (McdAB) system **(MacCready et al., 2018; MacCready et al., 2020; MacCready et al., 2021)**. Further suggesting that these puncta could represent assemblies of minimal wiffle balls, visualization of subcellular structures via TEM revealed the presence of multiple definable circular structures within the cytosol that appeared only in the cells induced to co-express HO BMC-H and BMC-T1 **(Figure 3C)**. No such circular structures were observable in control wildtype references **(Figure 3B)**. These circular bodies were 41.8 nm in diameter on average and resolved as a ‘clearance’ of other cytosolic structures lacking any electron-dense internal components **(Figure 3CD)**. These bodies are consistent with the defined minimal wiffle ball that has been assembled *in vitro* and structurally solved, with a diameter of 38 nm.

**Figure 3:**
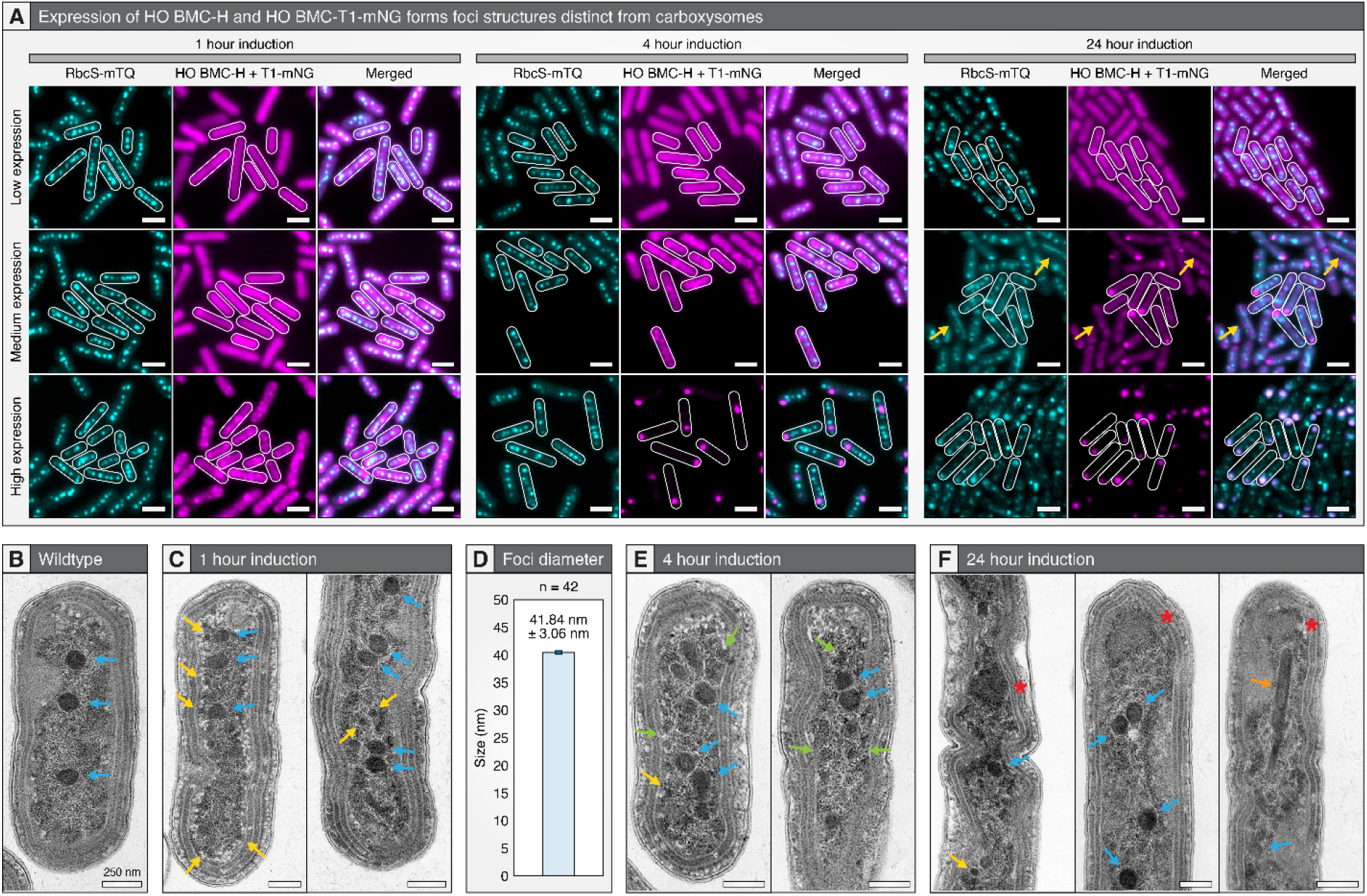
Expression duration influences resulting subcellular carboxysome and HO shell protein assembly ultrastructure. (A) Co-expressed HO BMC-H and T1-mNG displays localization patterns dependent on time and degree of induction. Low and/or short induction results in many distinct HO puncta separable from carboxysomes while high/long induction results in large peripheral puncta and disorganized carboxysome morphology. Scale bars = 2µm. (B) TEM micrograph of wildtype S. elongatus cell. (C) TEM micrograph at 1 hour of high induction. (D) Quantification of HO-like structures at 1 hour of high induction. (E) TEM micrograph at 4 hours of high induction. (F) TEM micrograph at 24 hours of high induction. Blue arrows indicate selected carboxysomes, yellow arrows indicate selected ∼40 nm circular structures only present in cells co-expressing HO BMC-H and T1 shell proteins. Other features evident at ≥4 hours after expression of HO BMC-H and HO BMC-T1 shell proteins include:, green arrows - indicate selected features with intermediate sizes between the ∼40nm foci and carboxysomes, and red asterisks indicate - mid-cell or polar localized amorphous electron dense structures, and orange arrow – apparent bar carboxysomes. TEM scale bars = 250 nm. Orange arrow—bar carboxysome, maybe HO hex aggregates with CcmL, so fewer for csome.

However, when expressed at moderate or high levels for a longer period of induction, HO BMC-H and HO-T1 no longer displayed a localization pattern consistent with individual, separable HO shell proteinfoci. At later time points (24 hours for medium induction and ≥ 4 hours for high), most HO shell proteins localized to a single, large punctum positioned at the pole of the cell. BMC aggregation is a common phenomenon in the absence of a positioning system **(Savage et al., 2010; MacCready et al., 2018; MacCready et al., 2021)**, yet at the resolution limits of light microscopy, it could not be determined if these represent aggregates of minimal wiffle balls. Furthermore, at the 24-hour time points for medium and high induction, substantial disruption of carboxysome morphology was evident at the resolution limits of light microscopy, including bar carboxysome formation, mispositioning, and “fragmentation” of the rubisco cargo signal **(Figure 3A – right panels)**.

TEM analysis of the corresponding samples revealed an unusually heterogeneous population of structures, most of which exhibited electron-dense interiors characteristic of the semi-crystalline packing of Rubisco in the lumen of carboxysomes. At 4 hours, cells contained both carboxysomes with relatively normal ultrastructural features, but also contained a subset of carboxysomes with perturbed morphology **(Figure 3E)**, an observation correlated with *in vivo* localization patterns of HO BMC-H to 1-3 carboxysomes at this timepoint **(Figure 2A)**. Moreover, a range of electron dense structures between the known sizes of HO shells (∼ 40 nm) and carboxysomes (∼ 175 nm) were also observed throughout the cytosol; we could not clearly classify these structures. At 24 hours, TEM visualized many cells with large intracellular aggregates, including bar carboxysomes **(Figure 3F - orange arrow)** and large electron dense structures frequently positioned near the mid-cell or at the cell pole **(Figure 3F – red asterisks)**.

We next swapped the mNG fluorescent reporter to HO BMC-H and left HO BMC-T1 untagged in order to visualize both HO components and observed similar results to the above. Previously, when HO BMC-H-mNG was expressed by itself, it colocalized to nearly all carboxysomes after 24 hours at low expression **(Figure 2A – right panel)**. However, co-expression of both HO BMC-H-mNG and HO BMC-T1 for 24 hours resulted in the formation of multiple small puncta and occasional small aggregates that did not colocalize with carboxysomes **(Supplemental figure S3A)**. Notably, there was no enrichment of HO BMC-H at carboxysomes when it was co-expressed with its cognate shell protein trimer, further supporting that HO BMC-T1 may preferentially nucleate HO BMC-H assembly and reduce its spurious inclusion into carboxysomes. However, we still observed a disruption of carboxysome morphology when HO BMC-H-mNG and BMC-T1 were co-expressed at a high level for an extended time period, indicative that co-expression could diminish, but not eliminate, cross-talk between the native carboxysome and the heterologous shell proteins. Taken together, the colocalization of Rubisco and HO BMC-H / HO BMC-T1 signals *in vivo* **(Figure 3A and Supplementary figure 3A)**, the absence of clear “edges” to the electron dense structures at polar regions of the cells **(Figure 3F)**, and the sequence similarity between HO BMC-H and carboxysome shell proteins **(Supplementary figure 1A)**, it seems probable that BMC HO BMC-H interferes with the maturation of procarboxysomes into mature carboxysomes. Iβ-carboxysome maturation is thought to initiate when CcmM nucleates Rubisco into phase-separated droplets, called procarboxysomes, that further recruit CcmK2 and other shell components via the adaptor protein CcmN **(Kinney et al., 2012, Wang et al., 2019; Zang et al., 2021)**. Two compatible hypotheses to explain these observations are: i) BMC HO BMC-H can integrate into carboxysome shells, interfering with normal facet assembly by interrupting normal self-assembly or blocking binding sites; or ii) HO BMC-H may recruit directly to the surface of Rubisco procarboxysomes via interaction with the CcmN encapsulation peptide. Both hypotheses could account for the intermediate sizes of electron dense structures we see at 4 hours and the large amorphous structures at 24 hours **(Figure 3EF)**.

### The McdAB positional system may possess moonlighting functions in maintaining BMC orthogonality

We recently identified and characterized a two-partner system, maintenance of carboxysome distribution A and B (McdAB), that are related to the broader ParAB family, and which are responsible for positioning β-carboxysomes in cyanobacteria and α-carboxysomes in proteobacteria **(Vecchiarelli et al., 2012; MacCready et al., 2018; MacCready et al., 2020; MacCready et al., 2021)**. Mechanistically, McdA proteins dimerize and nonspecifically bind the bacterial nucleoid in the presence of ATP **(MacCready et al., 2018)**. McdB, bound to carboxysomes, interacts with, and displaces McdA from the nucleoid by stimulating it’s inherent ATPase activity **(MacCready et al., 2018)**. The formation of localized McdA depletion zones around each McdB-bound carboxysomes results in the formation of concentration gradients of McdA along the nucleoid that individual carboxysomes utilize to position themselves the furthest away from their nearest neighbors **(MacCready et al., 2018)**. Importantly, recent evidence has suggested that a conserved C-terminal residue in McdB is important for mediating a direct physical connection to the shell proteins of cyanobacterial carboxysomes **(Basalla et al., 2023)**.

Since we observed that co-expression of HO BMC-H and HO BMC-T1 led to the appearance of foci in the short-term, but induced the formation of large peripherial aggregates upon longer expression, **(Figure 3A – right panels)** we sought to determine if the positional system could reposition and/or reduce formation of these polar bodies. We therefore directly tested if an orthogonal McdAB positioning system could operate in tandem with the native cyanobacterial carboxysome positioning to segregate putative HO shell assemblies. Because the α-McdAB system from *Halothiobacillus neapolitanus* is recently characterized, is evolutionarily distant from *S. elongatus*, and isolates as a monomer **(MacCready et al., 2021)**, we reasoned it would be a candidate positional system that could remain orthogonal to the native β-McdAB carboxysome system. We therefore designed a locus that expressed three copies of HO BMC-H and a single copy of HO BMC-T1 fluorescently fused at its c-terminus to mNG and α-McdB from *H. neapolitanus*. The partner α-McdA gene from *H. neapolitanus* was expressed as a separate element using the native promoter for *S. elongatus* β-McdA **(Supplemental Figure S4A)**. No disruption of native carboxysome positioning was observed even though α-McdA was constitutively expressed in our genetic design.

We found that introduction of the α-McdAB system prevented the formation of the large polar bodies when co-expressing HO BMC-H and BMC-T1, instead the signal localization for HO shell proteins was distributed in small foci throughout the cyanobacterial cytosol **(Figure 3A and Figure 4A)**. Even when the HO shell proteins were induced at high expression levels for a long period of time, the foci remained discreet and separable, suggestive that the introduced α-McdAB system could prevent the aggregation of HO shell foci **(Figure 4A)**. Notably, with α-McdAB, HO foci were also concentrated strongly along the medial axis of the cell in a pattern strongly resembling the cyanobacteria nucleoid **(Figure 4A and Video 1)**. Furthermore, the diffusion of the small HO puncta was constrained with α-McdAB, providing visual evidence of nucleoid tethering of these heterologous shell assemblies **(Video 1)**. Generally, the puncta of BMC HO BMC-H and HO BMC-T1-α-McdB distributed across the cell nucleoid without disrupting carboxysomal position, however, when HO shell proteins were induced with high expression and a longer induction time, some disordering or “clumping” of carboxysomes on the nucleoid was observed **(Figure 4A; high expression, 24 hours induction)**.

**Figure 4:**
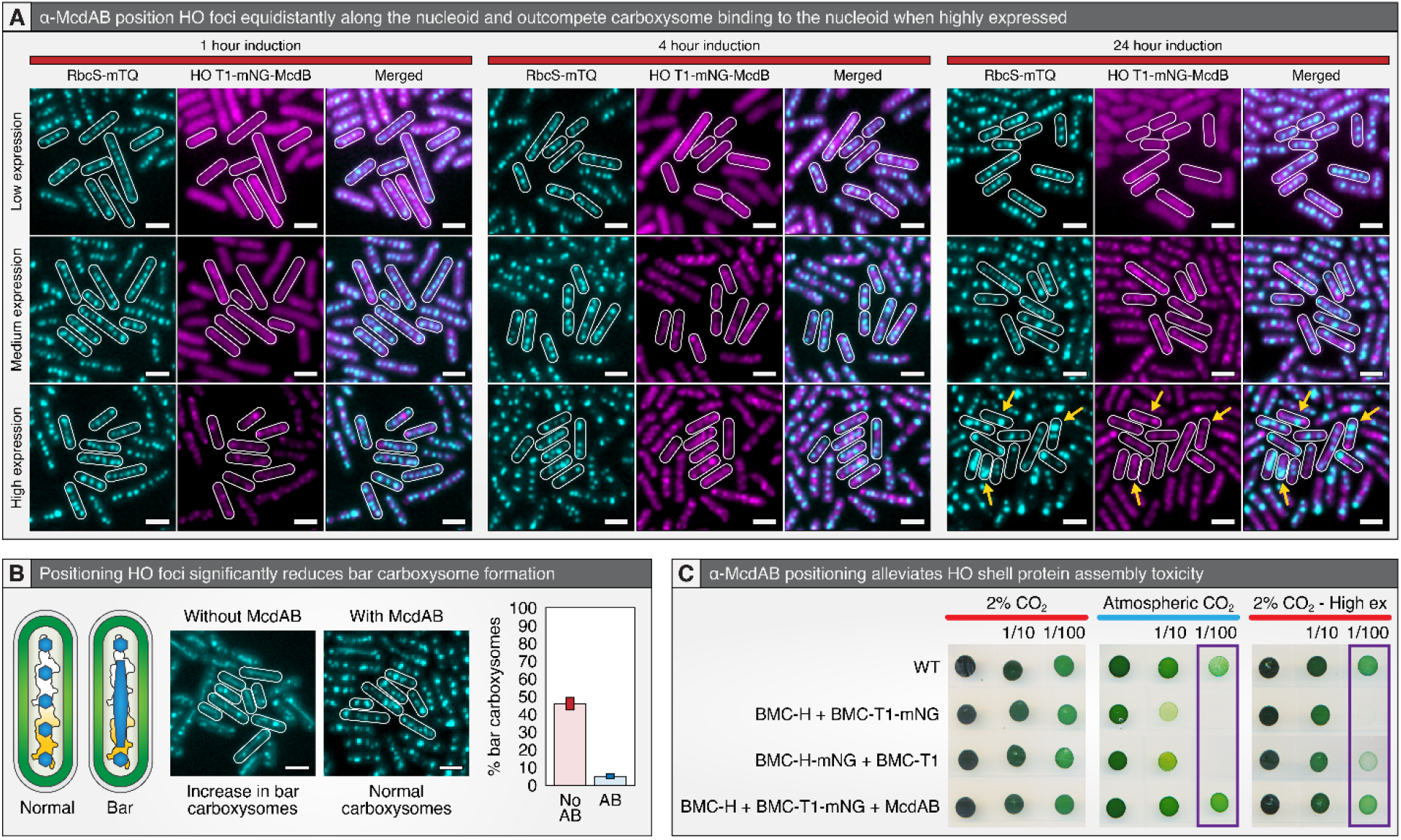
Orthogonal McdAB system prevents colocalization of HO and carboxysome shell components and rescues atmospheric CO_2_ growth impairment. (A) Co expression of HO BMC-H and T1-mNG-McdB. Scale bars = 2 µm. (B) Quantification of bar carboxysomes with and without the orthogonal McdAB system. (C) Serial diluted spot plate assay with HO and carboxysome shell proteins at 2% and atmospheric CO_2_ levels.

Unexpectedly, use of the heterologous α-McdAB positional system for the HO shell foci appeared to strongly reduce the pleotropic effects on the morphology of carboxysomes across all induction levels **(Figure 4A)**. We quantified carboxysome disruption as a function of the presence of bar carboxysomes (a visible indicator of severe carboxysome disorganization resolvable with fluorescence microscopy) **(Chen et al., 2013)**, finding that the the α-McdAB system significantly decreased bar carboxysome formation relative to a paired control lacking the heterologous positional system **(Figure 4B)**. Confirming this interpretation, we observed that including the α-McdAB positional system alleviated the growth inhibition of *S. elongatus* when the HO minimal wiffle ball constructs (i.e., HO BMC-H and HO BMC-T1) were induced: both with atmospheric CO_2_ and at high levels of HO expression under enriched CO_2_ headspace **(Figure 4C)**. As noted above, some clustering of carboxysomes was observed in the presence of a highly induced HO cassette at late time points, however, this phenotype was qualitatively different when the α-McdAB system was included **(Figure 3A vs. Figure 4A)** and the clustered carboxysomes appeared fully functional as assayed by cellular growth at atmospheric CO_2_ **(Figure 4C)**. Phenotypes of “overcrowding” of other ParAB cargo (*e.g*., high copy plasmids) upon a limited nucleoid “surface” provide precedence for this observation **(Million-Weaver S and Camps M, 2014)**. Taken together, we find that a heterologous α-McdAB positional system is not only able to tether the HO shell protein assemblies to the cyanobacterial nucleoid in a distributed fashion, but it also reduces the pleotropic growth effects and carboxysome morphology defects caused by HO BMC-H cross-reactivity.

Collectively, our results show for the first time that there can be considerable clash between the shell proteins of even evolutionarily-distant BMC types that are co-expressed in the same cell unless certain criteria are met. Multiple results suggest that some BMC shell proteins lack inherent orthogonality and are likely to cross-react with a broader array of shell proteins outside of their class **(*e.g*., HO BMC-H, Figure 2A)**. This cross-reactivity is apparent despite relatively large evolutionary distance between the carboxysome and HO BMC classes **(Figure 1D) (Melnicki et al., 2021; Sutter et al., 2022)**. A number of factors appear to assist in maintaining a separate identity of shell protein assemblies from different BMC classes. First, we find that the expression level of shell proteins can strongly influence the degree of cross-reactivity between different BMC classes. At low expression, HO BMC-H shell protein still exhibits limited co-localization with the carboxysome, but the impact of this cross-reactivity on carboxysomal morphology and function is less severe **(Figures 2A and 3A)**. Second, co-expression of multiple shell components from a given BMC class may help to restrict cross-reactivity for individual shell proteins **(Figure 3A)**. Stated differently, even cross-reactive shell proteins, such as HO BMC-H, may still display bias towards assembly with other shell proteins they co-evolved with; when multiple shell proteins from the same system are present, they may selectively ‘titrate out’ free subunits, reducing the cross-reactivity between systems. Third, our data also shows that introduction of a HO-tethered McdAB system strongly reduces the cross-reactivity of HO BMC-H, perhaps indicative that this family of proteins could have moonlighting functions in insulating the shell proteins from different microcompartments from interacting with one another. We observe that inclusion of a heterologous α-McdAB system not only allows for positioning of HO shell proteins in cyanobacteria, but it also significantly decreases the negative impact of heterologously expressing foreign shell proteins in cyanobacteria **(Figure 4)**. Finally, our data suggests that the process of carboxysome maturation may be the most susceptible to interference from HO BMC-H and therefore temporal segregation of shell protein expression could contribute to orthogonality. We observe gross carboxysome morphology is maintained under conditions of co-expression of HO BMC-H for up to 24 hours with when cultures are incubated in the dark **(Figure 2D)**, and that HO BMC-H co-localizes first to only 1 or 2 carboxysomes at early time points **(Figure 2A)**. However, as carboxysome dynamics remain poorly-characterized (especially under dark conditions) we regard this final point as speculative.

The mechanism for the observed rescue of inter-class shell protein cross-reactivity via the inclusion of a heterologous α-McdAB positional system remains unknown. In this regard, we note that all characterized McdB proteins are intrinsically disordered and exhibit rapid phase separation capabilities dependent on pH **(MacCready et al., 2020; MacCready et al., 2021)**. Recent analysis also has shown that the disordered protein domain responsible for phase separation capabilities are separable from C-terminal features necessary for shell protein interaction **(Basalla et al., 2023)**. One speculative hypothesis is that McdB’s oligomerization via phase separation may assist in nucleating distinct shell protein assemblies through physical association in a similar manner to cargo-shell interactions **(Basalla et al., 2023)**, thereby helping maintain separate arrays of BMC types by shell protein trafficking. This hypothesis receives indirect support from our observation that bacteria encoding more than one BMC type often encode for McdAB, and sometimes for an “orphan” McdB with no neighboring McdA **(Supplemental Figure S5)**. It is important to note that this hypothetical “moonlighting” function for McdB in the maintenance of distinct BMC shell architectures would require preferential interaction of McdB with specific shell proteins, which is currently an open question in the field of BMC biology.

In conclusion, we find that HO shell proteins can be co-expressed along with carboxysomes in the model organism *S. elongatus*. Maintenance of distinct microcompartments likely depends on a combination of suitable expression levels of shell proteins, cooperative binding effects of multiple shell proteins within the same BMC class, and/or moonlighting functions of the McdAB “positional” system. Our results have implications for organisms that naturally encode many different BMC types within their genome, such as *M. halophilus* **(Sutter et al., 2021)**. Future research to further elucidate features predictive of orthogonality between structural proteins of BMCs will be an important element to realize the potential of this class of microcompartment for biotechnological applications.

### Materials and Methods Construct designs

All constructs in this study were generated using Gibson Assembly **(Gibson et al., 2009)** from synthetized dsDNA (IDT) and verified by sequencing. Constructs contained flanking DNA that ranged ∼500-1000 bp in length upstream and downstream of the targeted insertion site, neutral site 1 or 2, to promote homologous recombination into target genomic loci **(Clerico et al., 2007)**. All HO shell protein expression was controlled through the use of a synthetic riboswitch driven by the Ptrc promoter and lacking *lacI* **(Nakahira et al., 2013)**. The carboxysome reporter, *rbcs*::RbcS-mTQ, was inserted upstream of each HO shell protein inserted at neutral site 2. For the positional construct, 3 codon optimized copies of HO BMC-H were inserted before HO BMC-T1-mNG fused at its c-terminus with the α-McdB protein from *Halothiobacillus neapolitanus* C2. The α-McdA sequence from *H. neapolitanus* C2 was inserted upstream of HO BMC-T1-mNG-McdB and its expression controlled using the native promoter for the β-McdA gene from *S. elongatus* 7942 **(Supplemental figure S4)**. All plasmid sequences for the constructs reported herein are deposited with Addgene for distribution and for referencing of plasmid design.

### Culture conditions and transformations

All *S. elongatus* cultures were grown in 125 mL baffled flasks (Corning) containing 50 ml BG-11 medium (SIGMA) buffered with 1 g/L HEPES to pH 8.3. Flasks were cultured in a Multitron II (atrbiotech.com) incubation system with settings: 120 µmol m−2 s−1 light intensity, 32°C, 2% CO_2_, shaking at 130 RPM. Cloning of plasmids was performed in *E. coli* DH5α chemically competent cells (Invitrogen). All *S. elongatus* transformations were performed as previously described **(Clerico et al., 2007)**. Cells were plated on BG-11 agar with either 12.5 mg ml−1 kanamycin or 25 mg ml−1 spectinomycin (depending on the antibiotic associated with the transformation plasmid). Single colonies were picked into 96-well plates containing 300 μl of BG-11 with identical antibiotic concentrations. Wells of cultures containing positive genetic insertion were then transferred to 125mL baffled flasks containing 50mL BG-11 and subjected to increasing antibiotic concentrations until complete penetrance was verified. Cultures were verified for complete insertion via PCR and removed from antibiotics prior to the initiation of any reported experiment.

### Induction conditions

For all reported data indicating a variable gene expression from the tunable riboswitch element the following concentrations of inducer were added at time 0; low expression = 25 µM, medium expression = 200 µM, high expression = 1000 µM. For spot plate assays, 2 ml of cell cultures were pelleted at 5000xg for 30 s, resuspended in 20 µl fresh BG-11, serial diluted, and 10 µl spotted onto BG-11 agar plates containing the respective concentration of theophylline (low – 25 µM or high – 1000 µM).

### Phylogenetic inference

Alignments for shell protein sequences were performed using MAFFT 1.3.7 under the G-INS-I algorithm and BLOSUM62 scoring matrix. A phylogenetic tree was then estimated with maximum likelihood analyses using RAxML 8.2.11 under the LG+Gamma scoring model of amino acid substitution.

### Fluorescence microscopy

All live-cell microscopy was performed using exponentially growing cells. Two mL of culture was spun down at 5000xg for 30 s, resuspended in 50 µl of BG-11 and 2 µl transferred to a square 2% agarose + BG-11 pad on glass slides. All images were captured using a Zeiss Axio Observer A1 microscope (100x, 1.46NA) with an Axiocam 503 mono camera. Image analysis was performed using Fiji v 1.0.

### Transmission Electron Microscopy

Cyanobacterial cultures were grown to OD750 = 0.7 in BG-11, induced with 1 mM theophylline for the respective duration (1, 4, or 24 hours), pelleted by centrifugation at 5,000 g for 3 minutes, and fixed overnight at 4°C with 2.5% formaldehyde / 2.5% glutaraldehyde in 0.1 M sodium cacodylate buffer (pH 7.4). Samples were then suspended in 2% high-purity agarose beads and cut into ∼1 mm cubes. Following three washes with 0.1 M sodium cacodylate buffer, cells were suspended in 1% osmium tetroxide / 1.5% potassium ferrocyanide and incubated overnight at 4°C. After incubation, cells were washed with HPLC-quality H2O until clear. Cells were then suspended in 1% uranyl acetate and microwaved for 2 min using a MS-9000 Laboratory Microwave Oven (Electron Microscopy Science), decanted, and washed until clear. Cells were dehydrated in increasing acetone series (microwave 2 min) and then embedded in Spurr’s resin (25% increments for 10 min each at 25°C). A final overnight incubation at room temperature in Spurr’s resin was done, then cells were embedded in blocks which were polymerized by incubation at 60°C for three days. Thin sections of approximately 50 nm were obtained using an MYX ultramicrotome (RMC Products), post-stained with 1% uranyl acetate and Reynolds lead citrate and visualized on a JEOL 1400 Flash transmission electron microscope (JEOL) equipped with an Orius SC200-830 CCD camera (Gatan). Contrast correction was performed using Fiji v 1.0.

## Acknowledgements

We would like to thank Alicia Withrow at the Michigan State University Center for Advanced Microscopy for preparation and thin-sectioning of samples. This work was supported as part of the Center for Catalysis in Biomimetic Confinement, an Energy Frontier Research Center funded by the U.S. Department of Energy, Office of Science, Basic Energy Sciences under Award Number DE-SC0023395. MED was supported by the Office of Science of the US Department of Energy DE-FG02-91ER20021 and MSU AgBio Research.

## Author Contributions

J.S.M. and M.E.D. designed the research, performed research, analyzed data and wrote the paper. C.A.K. designed the research and wrote the paper. D.C.D. designed the research, analyzed data and wrote the paper.

## Figure Legends

**Supplementary Figure S1: Expression of HO BMC-H-mNG for 1 hour**

(A) Expression of HO BMC-H-mNG at 1 hour of induction. Cells exhibit 1-2 HO BMC-H foci that are localized separately from the carboxysome reporter at this time point. Scale bars = 2 µm.

**Supplementary Figure S2: Exploring behavior similarities among HO BMC and carboxysome shell proteins**

(A) A complete set of representative images of CcmK2-mNG, CcmK3-mNG, and CcmK4-mG when expressed at 3 induction levels and at 3 time points (expansion of data set shown in Figure 2B). Native shell protein reporters colocalize with carboxysomes signal (RbcS-mTQ) at varying degrees dependent on time and degree of induction. (B) Representative images of HO BMC-H-mNG colocalization to carboxysomes following a treatment regimen of 24 hours growth in the light followed by 24 hours incubation in the dark under otherwise identical conditions. Scale bars = 2 µm.

**Supplementary Figure S3: Exploring HO BMC-H-mNG localization when coexpressed with T1**

(A) Co-expression of HO BMC-H-mNG and T1 as a comparison to the reciprocal tagging experiment in Figure 3A. Scale bars = 2 µm.

**Supplementary Figure S4: HO BMC positional system design**

(A) Cartoon schematic of the α-carboxysome genes near native operon structure containing α-McdAB in *H. neapolitanus* and the derived engineered HO BMC α-McdAB positional system characterized in Figure 4.

**Supplementary Figure 5: Positional systems in bacteria coding for more than one BMC type**

(A) Genomic neighborhood diagrams of bacteria encoding both PDU and EUT BMC-related components (yellow) show the presence of McdA (red) and/or McdB (blue) for one BMC type, but not the other.

**Video1: α-McdAB equidistantly position carboxysomes and HO foci on the nucleoid**

Carboxysomes (RbcS-mTQ, blue, left) are positioned simultaneously on the nucleoid with HO foci (HO BMC-H + HO BMC-T1-mTQ-McdB, red, middle) as distinct units (merged, right).

